# Canine saliva is a source of interspecies antimicrobial resistance gene transfer

**DOI:** 10.1101/2022.03.07.483304

**Authors:** Adrienn Gréta Tóth, Imre Tóth, Bernadett Rózsa, Eszter Gabriella Kovács, Attila Dubecz, Árpád V. Patai, Tibor Németh, Selçuk Kaplan, László Makrai, Norbert Solymosi

**Affiliations:** Centre for Bioinformatics, University of Veterinary Medicine Budapest, 1078 Budapest, Hungary; Department of Operative Tecniques and Surgical Research, Faculty of Medicine, University of Debrecen, 4032 Debrecen, Hungary; Department of Thoracic Surgery, Borsod-Abauj-Zemplén County Hospital, and University Teaching Hospital, 3526 Miskolc, Hungary; Department of Small Animal Clinical Sciences, College of Veterinary Medicine, Tuskegee University, Tuskegee, AL 36088, USA; Department of Surgery, Paracelsus Medical University, 90419 Nuremberg, Germany; Department of Surgery, Transplantation and Gastroenterology, Semmelweis University, 1085 Budapest, Hungary; Interdisciplinary Gastroenterology Working Group, Semmelweis University, 1085 Budapest, Hungary; Department and Clinic of Surgery and Ophthalmology, University of Veterinary Medicine Budapest, 1078 Budapest, Hungary; Tekirdag Namik Kemal University, Faculty of Veterinary Medicine, Department of Genetics, 59100 Tekirdag, Turkey; Department of Microbiology and Infectious Diseases, University of Veterinary Medicine Budapest, Budapest, 1143 Hungary; Eötvös Loránd University, Department of Phyisics of Complex Systems, 1117 Budapest, Hungary

## Abstract

While the One Health issues of intensive animal farming are commonly discussed, keeping companion animals is less associated with the interspecies headway of antimicrobial resistance. With the constant advance of veterinary standards, antibiotics are regularly applied in companion animal medicine. Due to the close coexsistance of dogs and humans, dog bites and other casual encounters with dog saliva (e.g. licking the owner) are common. According to our metagenome studies based on 26 new generation sequencing canine saliva datasets from 2020 and 2021 reposited in NCBI SRA by The 10,000 Dog Genome Consortium and the Broad Institute within Darwin’s Ark project, canine saliva is rich in bacteria with predictably transferable antimicrobial resistance genes (ARGs). In the genom of potentially pathogenic *Bacteroides, Capnocytophaga, Corynebacterium, Fusobacterium, Pasteurella, Porphyromonas, Staphylococcus* and *Streptococcus* species, that are some of the most relevant bacteria in dog bite infections, ARGs against aminoglycosides, carbapenems, cephalosporins, glycylcyclines, lincosamides, macrolides, oxazolidinone, penams, phenicols, pleuromutilins, streptogramins, sulfonamides and tetracyclines could be identfied. Several ARGs, including ones against amoxicillin-clavulanate, the most commonly applied antibiotic by dog bites, was predicted to be potentially transferable based on their association with mobile genetic elements (e.g. plasmids, phages, integrated mobile genetic elements). According to our findings canine saliva may be a source of transfer of ARG-rich bacteria, that can either colonize the human body or transport ARGs to the host bacteriota and thus can be considered as a risk in the spread of antimicrobial resistance.

## Background

Antimicrobial resistance is a threat of utmost significance that constantly raises medical challenges all around the globe. The fact that main drivers of the headway of antimicrobial resistance (AMR) are antimicrobial use and abuse is well-accepted. Moreover, according to the concept of One Health, the effects of antimicrobial use in human, animal and environmental sectors are interconnected, and thus interdependent. In case of AMR, the relatedness of human and animal dimensions described by the One Health approach can be the best referred by considering that most antimicrobial classes are co-used in both sectors. Human health antimicrobial use has been overshadowed for years by farm animal mass medication, although this tendency has recently changed at some parts of the world^1^. While the appearance and advance of AMR, and as an underlying cause, the enrichment and transmission of antimicrobial resistance genes (ARGs) in antibiotic-dense environments, such as intensive animal production farms is a well-examined phenomenon^2,3^, the spread of AMR may also derive from other animal-borne routes.

Over the past decades, the number of companion animals has been tendentially and steadily rising^4^. Between 2000 and 2017 the number of dogs in the United States escalated from 68 million to 89.7 million^5^. 67.9% of all households in the U.S. was associated with the ownership of various pet species and 48% of all with dogs in 2016^4^. In years 2019–2020 50% of U.S. population owned a dog^6^. Coronavirus Disease (COVID-19) Pandemic outbreak has resulted in elevated companion animal acquisition rates, albeit often followed by retention or replacement^7,8^. Besides the popularity of keeping small animals, the quality of human-pet bond has also changed. According to the survey of the American Veterinary Medical Association, 70% of pet owners consider their pets as family members, 17% as companions and 3% as property^9^. The role of pets can principally be defined as social companionship. Nowadays having physical proximity is very common by pet-owner coexistences, pets often sleep together with their owners and lick their face or wounds^10^. Unfortunately and unsurprisingly, with such high dog numbers, the occurrence of dog bites is also common. Between 2001 and 2003, approximately 4,5 million dog bites were registered yearly in the United States, 19% of which necessitated medical intervention^11^. In years 2005–2013 an average of 337,103 dog bite injuries were treated at the U.S. emergency departments^12^, although dog bites in general are under-reported^13^. Interestingly, 3 of 5 bites are executed by family dogs, what is more common than attacks by strays^14^. According to some statistics, some dog breeds including pit bulls, rottweilers and German shepherds are more likely to commit attacks, although any dog breeds may bite^15^. According to the current ranking of the American Kennel Club, German shepherds are the 3rd and rottweilers are the 8th most popular dog breeds, while pit bulls are banned in many states.

According to a publication from 2017, dogs make up 64.8% of the veterinary-visiting population in Great Britan^16^. Another study executed on a representative number of U.S. residents suggests that approximately 90% of cat and dog owners had taken a visit at a veterinarian at any time and about 40% visited a veterinarian annually^17^.

The modern mindset of providing regular veterinary healthcare services to our pets and keeping them in our closest surroundings may contribute to the interspecies transmission of AMR. Several studies have already turned the attention to the role of companion animals in the headway of AMR^18–22^. Nevertheless, the significance of the direct pet-borne AMR spread route has been given less attention when compared with the rather indirect, mostly food-transmitted farm animal associated route. The routes of ARG transmission can be assessed by analyzing the genetic surroundings of ARGs. Genetic elements that facilitate horizontal gene transfer (HGT) including plasmids, bacteriophages and integrative mobile genetic elements (iMGEs) may contribute to three different HGT processes, namely conjugation, transduction or transformation. By conjugation, genes are delivered to a recipient cell on plasmids if bacterial cells are physically binded, while by transduction, direct cell-to-cell contact is disclaimed due to the presence of bacteriophages, as means of gene conduit. Transformation negates the need for particular delivery processes. In this case, bacteria take up genetic fragments from their environment^23^. After dog bites or close encounters with saliva from dogs that undergo veterinary treatments and thus carry bacteria with an enriched ARG content, resistant bacteria may be introduced to the human body and later the HGT of antimicrobial resistance determinants may be exchanged with the host bacteriota. Our study aimed to reveal the ARG content of 26 canine saliva samples from the U.S., attach the ARGs with the bacterial species that they derive from and report the ARGs’ spreading capabilities to weigh the above-mentioned phenomenon. For this purpose, freely accessible next-generation sequencing (NGS) shotgun datasets were downloaded and bioinformatically analyzed using an advanced metagenomic pipeline.

## Materials and Methods

Deep sequenced canine saliva datasets were searched in the National Center for Biotechnology Information (NCBI) Sequence Read Archive (SRA) repository. In May 2021, two shotgun metagenomic BioProjects (PRJNA64812377 – The 10,000 Dog Genome Consortium, PRJNA683923 – Broad Institute, Darwin’s Ark project) with more than 100,000,000 paired-end reads per sample were identified (Table 1). The median read count (interquartile range, IQR) of the samples was 177.7 × 10^6^ (26.6 × 10^6^) and 417.7 × 10^6^ (90.1 × 10^6^) in datasets PRJNA648123 and PRJNA683923, respectively.

**Table 1.** The list of analyzed samples obtained from National Center for Biotechnology Information Sequence Read Archive. Column Run contains the NCBI SRA run identifiers. Bacterial read count represents the number of the reads were classified taxonomically to any bacteria.

| ID | BioProject | Run | Sex | Bacterial read count |
| --- | --- | --- | --- | --- |
| 1 | PRJNA648123 | SRR12330029 | female | 2,900,387 |
| 2 |  | SRR12330041 | female | 16,153,172 |
| 3 |  | SRR12330042 | male | 13,072,781 |
| 4 |  | SRR12330043 | male | 13,774,332 |
| 5 |  | SRR12330044 | male | 6,123,646 |
| 6 |  | SRR12330045 | female | 16,707,766 |
| 7 |  | SRR12330098 | female | 18,826,266 |
| 8 |  | SRR12330104 | male | 27,598,592 |
| 9 |  | SRR12330220 | male | 9,938,948 |
| 10 |  | SRR12330260 | female | 17,642,933 |
| 11 |  | SRR12330298 | female | 17,277,697 |
| 12 |  | SRR12330356 | male | 13,988,719 |
| 13 |  | SRR12330364 | female | 17,378,513 |
| 14 |  | SRR12330377 | male | 12,155,726 |
| 15 |  | SRR12330378 | female | 34,183,357 |
| 16 |  | SRR12330382 | female | 22,353,314 |
| 17 |  | SRR12330383 | male | 22,886,951 |
| 18 |  | SRR12330384 | male | 18,328,656 |
| 19 |  | SRR12330385 | female | 6,631,504 |
| 20 | PRJNA683923 | SRR13340534 | female | 0 |
| 21 |  | SRR13340535 | female | 6,752,169 |
| 22 |  | SRR13340537 | male | 8,245,374 |
| 23 |  | SRR13340538 | male | 41,212,470 |
| 24 |  | SRR13340539 | female | 13,028,655 |
| 25 |  | SRR13340540 | female | 6,964,460 |
| 26 |  | SRR13340541 | male | 6,279,921 |

### Bioinformatic analysis

Quality based filtering and trimming of the raw short reads was performed by TrimGalore (v.0.6.6, https://github.com/FelixKrueger/TrimGalore), setting 20 as a quality threshold. Only reads longer than 50 bp were retained and taxonomically classified using Kraken2 (v2.1.1)^24^ and a database created (24/03/2021) from the NCBI RefSeq complete archaeal, bacterial, viral and plant genomes. For this taxon assignment the –confidence 0.5 parameter was used to obtain more precise species level hits. The taxon classification data was managed in R^25^ using functions of the packages phyloseq^26^ and microbiome.^27^ Reads were classified as origin of bacteria were assembled to contigs by MEGAHIT (v1.2.9)^28^ using default settings. The contigs were also classified taxonomically by Kraken2 with the same database as above. From the contigs all possible open reading frames (ORFs) were gathered by Prodigal (v2.6.3)^29^. The protein translated ORFs were aligned to the ARG sequences of the Comprehensive Antibiotic Resistance Database (CARD, v.3.1.3)^30,31^ by Resistance Gene Identifier (RGI, v5.2.0) with Diamond^32^ The ORFs classified as perfect or strict were further filtered with 90% identity and 90% coverage. All nudged hits were excluded. The integrative mobile genetic element (iMGE) content of the ARG harbouring contigs was analyzed by MobileElementFinder (v1.0.3) and its database (v1.0.2).^33^ Following the distance concept of Johansson et al.^33^ for each bacterial species, only those with a distance threshold defined within iMGEs and ARGs were considered associated. In the MobileElementFinder database (v1.0.2) for *Bacteroides*, the longest composite transposon (cTn) was the *Tn6186*. In case of this genus, its length (8,505 bp) was taken as the cut-off value. For the genera *Enterococcus* and *Klebsiella* the *Tn6246* (5,147 bp) and *Tn125* (10,098 bp) provided the threshold, respectively. In the case of *E. coli*, this limit was the length of the *Tn1681* transposon, namely 24,488 bp, while for *P. aeruginosa Tn6060* (25,440 bp). As the database neither contains species-level, nor genus-level cTn data for the rest of the species, a general cut-off value was chosen for the contigs of these species. This value was declared as the median of the longest cTns per species in the database (10,098 bp).

The plasmid origin probability of the contigs was estimated by PlasFlow (v.1.1)^34^ The phage content of the assembled contigs was prediced by VirSorter2 (v2.2.3)^35^. The findings were filtered for dsDNAphages and ssDNAs. All data management procedures, analyses and plottings were performed in R environment (v4.1.0).^25^

## Results

After presenting the bacteriome and the identified AGRs (resistome), predictions regarding the mobility potential of ARGs were also resumed based on genetic characteristics that may play a significant role in HGT.

### Bacteriome

By taxon classification, the number of reads aligning to bacterial genomes differed in the various samples. In the saliva median bacterial read count of the samples was 4.3 × 10^6^ (IQR: 3.4 × 10^6^).

The relative abundances of genera that achieved more than 1% of the bacterial hits in any of the saliva samples is shown in Figure 1. In the saliva samples the dominant genera (with mean prevalence) in descending order were *Porphyromonas* (49%), *Prevotella* (15%), *Pasteurella* (12%), *Neisseria* (10%), *Capnocytophaga* (9%), *Conchiformibius* (7%), *Frederiksenia* (7%), *Cutibacterium* (6%), *Actinomyces* (5%), *Campylobacter* (4%), *Desulfomicrobium* (4%), *Bacteroides* (3%), *Fusobacterium* (3%), *Mycoplasmopsis* (3%), *Treponema* (3%), *Streptococcus* (2%). In the sample No. 20 no reads were classified to bacteria.

**Figure 1.**
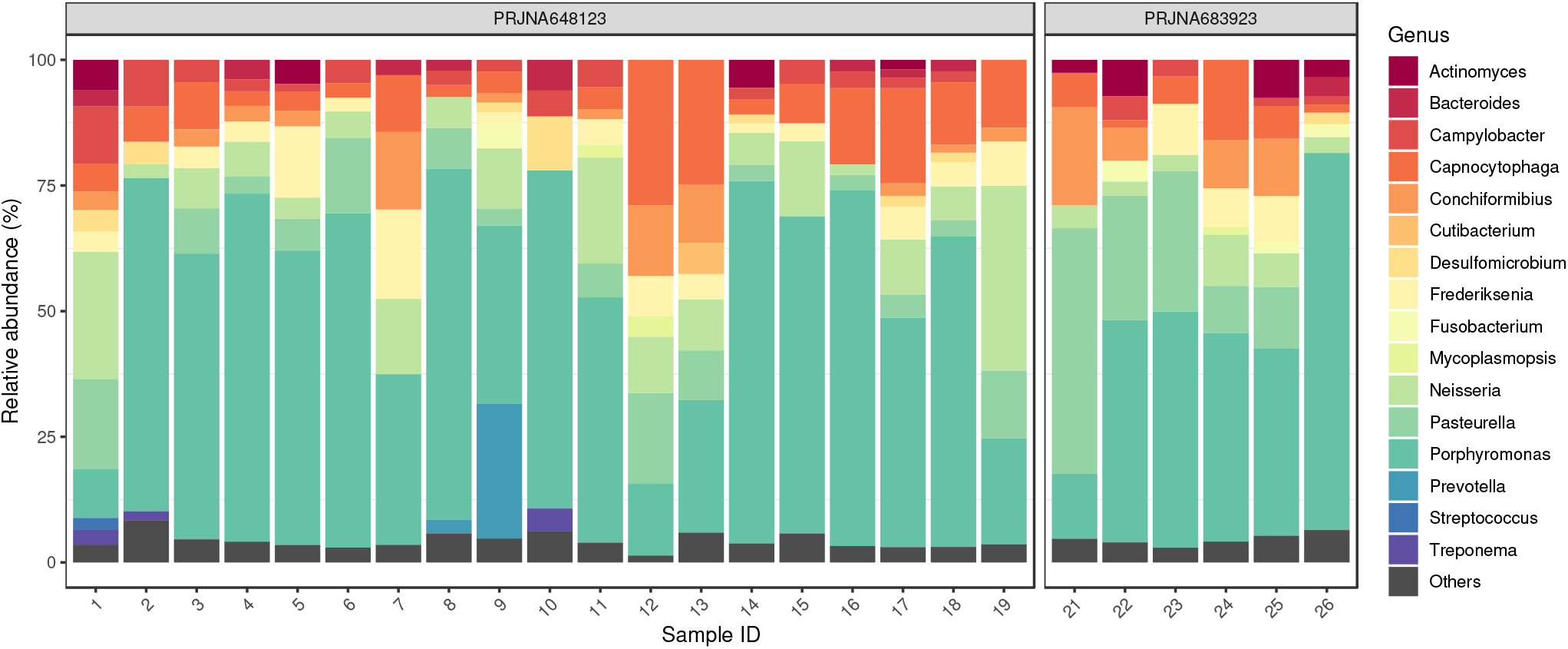
Saliva core bacteriome. The relative abundances of genera that achieved more than 1% of the bacterial hits in any of the samples. The dominant genera (with mean prevalence) in descending order were *Porphyromonas* (49%), *Prevotella* (15%), *Pasteurella* (12%), *Neisseria* (10%), *Capnocytophaga* (9%), *Conchiformibius* (7%), *Frederiksenia* (7%), *Cutibacterium* (6%), *Actinomyces* (5%), *Campylobacter* (4%), *Desulfomicrobium* (4%), *Bacteroides* (3%), *Fusobacterium* (3%), *Mycoplasmopsis* (3%), *Treponema* (3%), *Streptococcus* (2%). In the sample No. 20 no reads were classified to bacteria.

### Resistome

The dominant mechanism of identified ARGs was the antibiotic inactivation (49.44%), antibiotic target protection (22.63%), antibiotic target alteration (15.36%), antibiotic efflux (7.54%), antibiotic target replacement (5.03%).

The detected bacterial species were associated with the following ARGs: *Acinetobacter baumannii: aadA2, OXA-2; Actinobacillus pleuropneumoniae: ROB-1; Aeromonas hydrophila: ant(3”)-IIa, OXA-2, sul1; Alistipes indistinctus: tetQ; A. shahii: ermF, OXA-347; Amedibacterium intestinale: ant(6)-Ib, tet(44); Bacteroides dorei: tetQ; B. fragilis: aadS, ermF, mef(En2), OXA-347, tet(X5), tetX; B. heparinolyticus: OXA-347, tetQ; B. ovatus: tetQ; Bacteroides* sp. HF-5287: *tetQ; Bibersteinia trehalosi*: *ROB-10*; *Blautia hansenii*: *tet32*; *Bulleidia* sp. zg-1006: *tet32*; *Capnocytophaga cynodegmi*: *cfxA2*; *Capnocytophaga* sp. H2931: *aadS, OXA-347; Capnocytophaga* sp. H4358: *aadS, OXA-347; C. stomatis: aadS, ermF, OXA-347*, *tet(X5)*; *Chryseobacterium indologenes*: *aadS*, *ermF*; *Chryseobacterium* sp. POL2: *OXA-347*; *Clostridioides difficile: aac(6’)-Im, aph(2”)-IIa, ermG, tet32, tetM, tetO; Clostridium cellulovorans: tet32; Conchiformibius steedae: ROB-1; Corynebacterium* sp. 1959: *aph(3”)-Ib, aph(3’)-Ia, aph(6)-Id, sul2; Elizabethkingia anophelis: OXA-347; Empedobacter brevis: OXA-347; Enterocloster bolteae: tetW*; *Enterococcus faecalis: tetM; E.faecium: aad(6), aph(3’)-IIIa; E. gilvus: ermB; E. hirae: tetO; Enterococcus* sp. FDAARGOS_375: *ermB; Escherichia coli: acrA, aph(3’)-Ia, aph(3’)-IIa, emrE, emrK, gadW, gadX, mdtN, TEM-116, ugd; Eubacterium maltosivorans: tet32; Eubacterium* sp. NSJ-61: *tet32; Faecalibacterium prausnitzii: tet32, tetW*; *Filifactor alocis: tet(W/N/W); Fusobacterium ulcerans: OXA-85; Geobacter daltonii: ereA; G. sulfurreducens: OXA-119; Glaesserella parasuis: ROB-1, ROB-9; Haemophilus haemolyticus: ROB-1; Ha. parahaemolyticus: aph(3”)-Ib, sul2; Klebsiella michiganensis: aph(3”)-Ib, aph(6)-Id, catIII, dfrA14, sul2; K. quasipneumoniae: aph(3’)-Ia; Lachnoanaerobaculum umeaense: tet32; Megasphaera stantonii: tetW*; *Mogibacterium pumilum: tet(W/N/W), tetM; Moraxella bovis: aph(3”)-Ib, sul2; Murdochiella vaginalis: tetO; Myroides odoratimimus: OXA-347; Neisseria animaloris: aadA3; N. shayeganii*: *aph(6)-Id*; *Ochrobactrum anthropi*: *dfrA14*; *Parabacteroides distasonis*: *cfxA2*, *ermF*, *tetX*; *Pasteurella multocida*: *sul2; Peptoclostridium acidaminophilum: tet32; Phocaeicola coprophilus: tetQ; Porphyromonas cangingivalis: cfxA2, ermF, mef(En2)*; *P. crevioricanis*: *cfxA2*, *tetQ*; *P. gingivalis*: *cfxA2*, *mef(En2)*, *pgpB*; *Prevotella fusca*: *tetQ*; *P. intermedia*: *ermF*, *mef(En2)*, *tetQ*, *tetX*; *Proteus vulgaris*: *tet(H)*; *Providencia rettgeri*: *aph(6)-Id*, *sul2*; *Pseudomonas aeruginosa*: *OXA-2*, *qacL, sul1; P. putida: tet(H); Ralstonia insidiosa: OXA-573; R. pickettii: OXA-22, OXA-60; Riemerella anatipestifer: aadS, ermF, OXA-347, tet(X4), tet(X5), tetX; Roseburia intestinalis: tet32; Rothia nasimurium: tet(Z); Staphylococcus aureus: aad(6), aph(3’)-IIIa, SAT-4; Streptococcus acidominimus: tetO; S. agalactiae: aph(3’)-IIIa, ermB; S. anginosus: tet32, tetO; S. constellatus: tet32, tetO; S. equi: lnuC, tet32, tetO; S. gwangjuense: lnuC; S. parauberis: tetS; S. pluranimalium: mel; Streptococcus* sp. FDAARGOS_521: *tetM; S. suis: ermB, lnuB, lsaE, tetO, tetW*; *Tannerella forsythia: cfxA2; Trueperella pyogenes: ermX; Variovorax* sp. PAMC28562: *aph(3’)-Ia; Variovorax* sp. SRS16: *aph(3”)-Ib, aph(6)-Id*.

**Figure 2.**
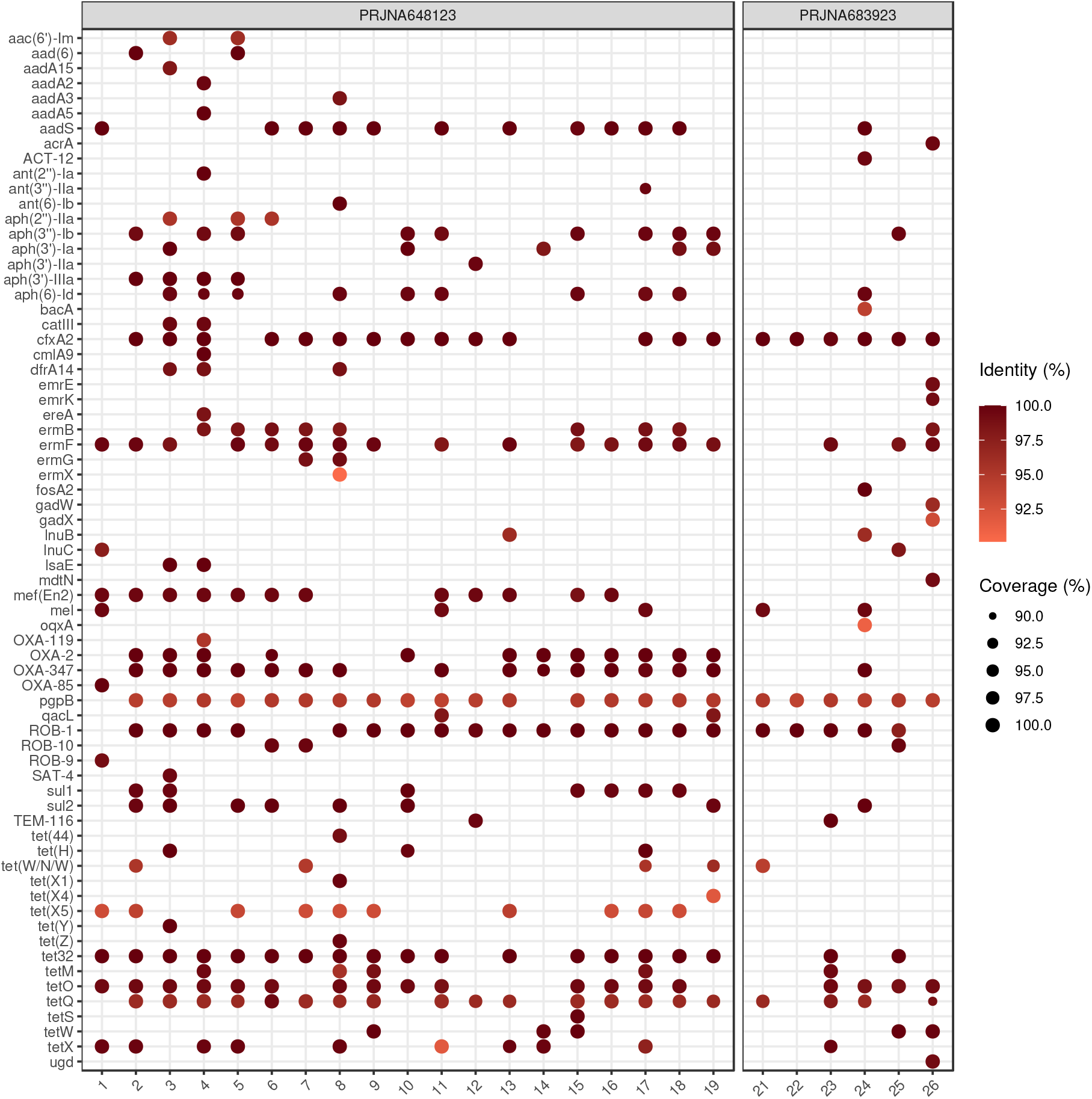
Identifed antimicrobial resistance genes (ARGs) by samples. For each sample-ARG combination, only the best finding is plotted. The size and the colour of the dots correspond to the coverage and the sequence identity of hits on reference genes, respectively. In sample No. 20 there was no identifiable ARG. The gene names that are too long have been abbreviated (*acrA: Escherichia coli acrA; emrE: E. coli emrE*).

Based on the ARG content, the detected bacterial species may show resistance against the following antibiotic groups: *Acinetobacter baumannii:* aminoglycoside, carbapenem, cephalosporin, penam; *Actinobacillus pleuropneumoniae:* cephalosporin, penam; *Aeromonas hydrophila*: aminoglycoside, carbapenem, cephalosporin, penam, sulfonamide; *Alistipes indistinctus*: tetracycline; *A. shahii:* carbapenem, cephalosporin, lincosamide, macrolide, penam, streptogramin; *Amedibacterium intestinale*: aminoglycoside, tetracycline; *Bacteroides dorei*: tetracycline; *B. fragilis*: aminoglycoside, carbapenem, cephalosporin, glycylcycline, lincosamide, macrolide, penam, streptogramin, tetracycline; *B. heparinolyticus*: carbapenem, cephalosporin, penam, tetracycline; *B. ovatus*: tetracycline; *Bacteroides* sp. HF-5287: tetracycline; *Bibersteinia trehalosi*: cephalosporin, penam; *Blautia hansenii:* tetracycline; *Bulleidia sp. zg-1006:* tetracycline; *Capnocytophaga cynodegmi:* cephamycin; *Capnocytophaga* sp. H2931: aminoglycoside, carbapenem, cephalosporin, penam; *Capnocytophaga* sp. H4358: aminoglycoside, carbapenem, cephalosporin, penam; *C. stomatis*: aminoglycoside, carbapenem, cephalosporin, lincosamide, macrolide, penam, streptogramin, tetracycline; *Chryseobacterium indologenes*: aminoglycoside, lincosamide, macrolide, streptogramin; *Chryseobacterium* sp. POL2: carbapenem, cephalosporin, penam; *Clostridioides difficile:* aminoglycoside, lincosamide, macrolide, streptogramin, tetracycline; *Clostridium cellulovorans*: tetracycline; *Conchiformibius steedae*: cephalosporin, penam; *Corynebacterium* sp. 1959: aminoglycoside, sulfonamide; *Elizabethkingia anophelis*: carbapenem, cephalosporin, penam; *Empedobacter brevis:* carbapenem, cephalosporin, penam; *Enterocloster bolteae:* tetracycline; *Enterococcus faecalis:* tetracycline; *E.faecium:* aminoglycoside; *E. gilvus:* lincosamide, macrolide, streptogramin; *E. hirae:* tetracycline; *Enterococcus* sp. FDAARGOS_375: lincosamide, macrolide, streptogramin; *Escherichia coli:* acridine dye, aminoglycoside, cephalosporin, disinfecting agents and intercalating dyes, fluoroquinolone, glycylcycline, macrolide, monobactam, nucleoside, penam, penem, peptide, phenicol, rifamycin, tetracycline, triclosan; *Eubacterium maltosivorans*: tetracycline; *Eubacterium* sp. NSJ-61: tetracycline; *Faecalibacterium prausnitzii:* tetracycline; *Filifactor alocis:* tetracycline; *Fusobacterium ulcerans:* carbapenem, cephalosporin, penam; *Geobacter daltonii:* macrolide; *G. sulfurreducens:* carbapenem, cephalosporin, penam; *Glaesserella parasuis:* cephalosporin, penam; *Haemophilus haemolyticus:* cephalosporin, penam; *H. parahaemolyticus:* aminoglycoside, sulfonamide; *Klebsiella michiganensis*: aminoglycoside, diaminopyrimidine, phenicol, sulfonamide; *K. quasipneumoniae*: aminoglycoside; *Lachnoanaerobaculum umeaense*: tetracycline; *Megasphaera stantonii*: tetracycline; *Mogibacterium pumilum*: tetracycline; *Moraxella bovis*: aminoglycoside, sulfonamide; *Murdochiella vaginalis*: tetracycline; *Myroides odoratimimus*: carbapenem, cephalosporin, penam; *Neisseria animaloris:* aminoglycoside; *N. shayeganii:* aminoglycoside; *Ochrobactrum anthropi:* diaminopyrimidine; *Parabacteroides distasonis:* cephamycin, glycylcycline, lincosamide, macrolide, streptogramin, tetracycline; *Pasteurella multocida:* sulfonamide; *Peptoclostridium acidaminophilum:* tetracycline; *Phocaeicola coprophilus:* tetracycline; *Porphyromonas cangingivalis:* cephamycin, lincosamide, macrolide, streptogramin; *P. crevioricanis:* cephamycin, tetracycline; *P. gingivalis*: cephamycin, macrolide, peptide; *Prevotella fusca*: tetracycline; *P. intermedia*: glycylcycline, lincosamide, macrolide, streptogramin, tetracycline; *Proteus vulgaris*: tetracycline; *Providencia rettgeri*: aminoglycoside, sulfonamide; *Pseudomonas aeruginosa:* carbapenem, cephalosporin, disinfecting agents and intercalating dyes, penam, sulfonamide; *P. putida*: tetracycline; *Ralstonia insidiosa*: carbapenem, cephalosporin, penam; *R. pickettii*: carbapenem, cephalosporin, penam; *Riemerella anatipestifer*: aminoglycoside, carbapenem, cephalosporin, glycylcycline, lincosamide, macrolide, penam, streptogramin, tetracycline; *Roseburia intestinalis*: tetracycline; *Rothia nasimurium*: tetracycline; *Staphylococcus aureus*: aminoglycoside, nucleoside; *Streptococcus acidominimus:* tetracycline; *S. agalactiae:* aminoglycoside, lincosamide, macrolide, streptogramin; *S. anginosus*: tetracycline; *S. constellatus*: tetracycline; *S. equi*: lincosamide, tetracycline; *S. gwangjuense*: lincosamide; *S. parauberis:* tetracycline; *S. pluranimalium:* lincosamide, macrolide, oxazolidinone, phenicol, pleuromutilin, streptogramin, tetracycline; *Streptococcus* sp. FDAARGOS_521: tetracycline; *S. suis*: lincosamide, macrolide, oxazolidinone, phenicol, pleuromutilin, streptogramin, tetracycline; *Tannerella forsythia:* cephamycin; *Trueperella pyogenes:* lincosamide, macrolide, streptogramin; *Variovorax* sp. PAMC28562: aminoglycoside; *Variovorax* sp. SRS16: aminoglycoside

### Mobilome

The frequencies of iMGEs, phages and plasmids associated with ARGs by bacteria of origin are summarized in Figure 3.

**Figure 3.**
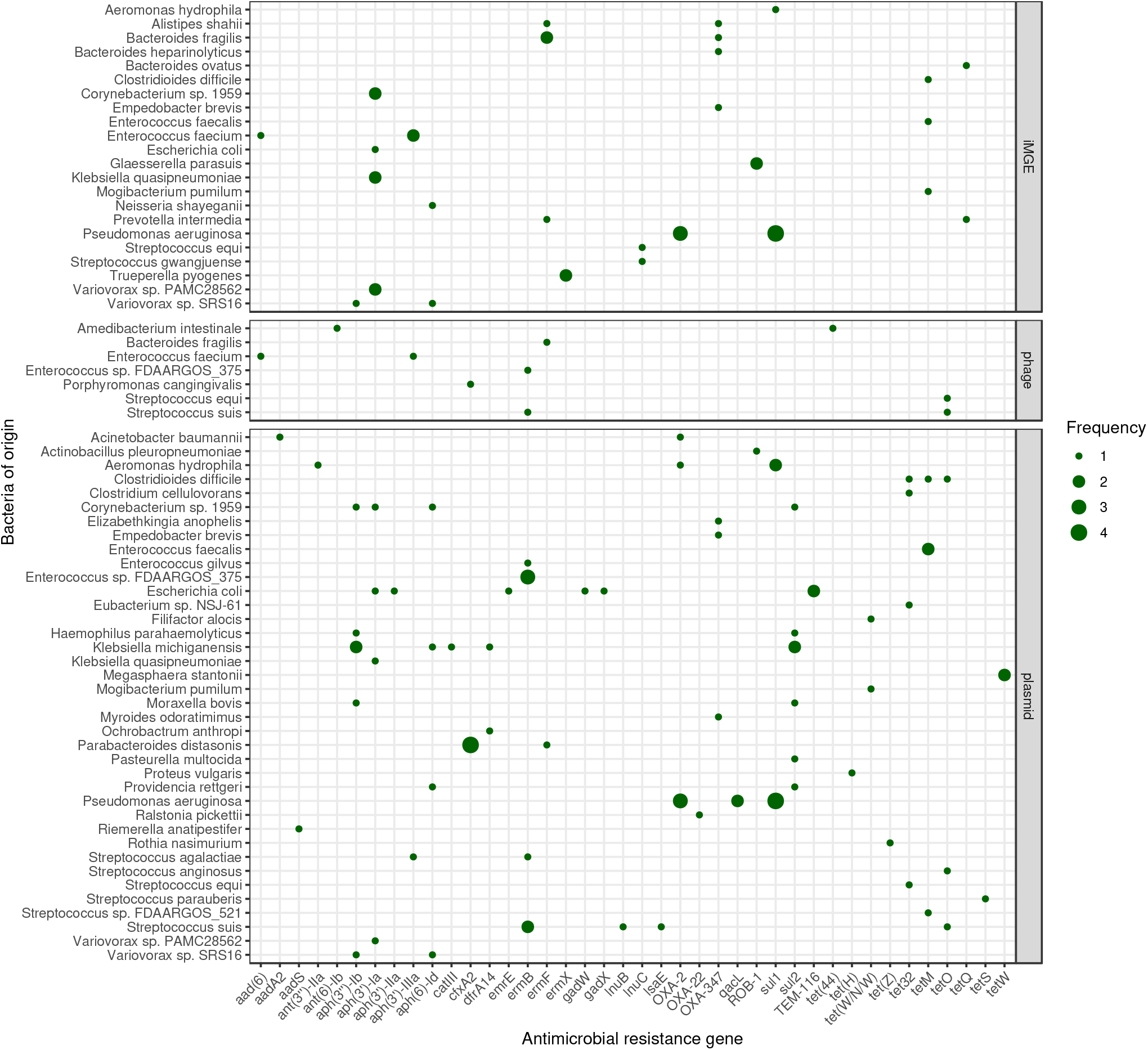
Mobile antimicrobial resistance gene frequency by bacteria of origin. The size of the dots indicates the occurrence frequency of the given gene flanked by iMGE, positioned in plasmid or phage.

## Discussion

During the bacteriome, resistome and mobilome analysis of the canine saliva samples a large set of results was obtained that can be examined from a one health point of view, merging the small animal veterinary sector with the perspective of the human healthcare system.

A total of 16 major bacterial genera were detected within the saliva samples, out of which several aerobic and anaerobic genera often get isolated from infected dog bite wounds. Dog bite infections are normally polymicrobial and the bite wound bacteriota consists of bacteria from the animals’ oral cavity, the recipients’ skin and the environment. The most common pathogens in dog bites are *Pasteurella* spp. (P. *canis* and P. *multocida), Staphylococcus* spp., *Streptococcus* spp. and *Capnocytophaga* spp., *Porphyromonas* spp., *Bacteroides* spp., *Fusobacterium* spp. and *Corynebacterium* spp.^36^ that all appeared in the analyzed saliva samples. Some other bacterial groups of a relatively higher clinical significance that were detected in the saliva samples including *Enterococcus* spp., *Moraxella* spp., *Neisseria* spp., *Prevotella* spp., *Pseudomonas* spp. are also often isolated from dog bite wound infections. The vast majority of other genera isolated in the samples have been mentioned to appear in dog saliva in previous publications with variable abundance rates^13,37^. Even though members of *Clostridium* spp. were detected in the samples, genome fragments of *C. tetani*, the bacterium responsible for tetanus were not identified.

The number of detected ARGs was relatively high in the salivary bacteriom. Examining 8 genera (*Pasteurella* spp., *Staphylococcus* spp., *Streptococcus* spp., *Capnocytophaga* spp., *Porphyromonas* spp., *Bacteroides* spp., *Fusobacterium* spp. and *Corynebacterium* spp.) that were indicated to be the most relevant ones in dog bite infections by other authors^36,37^ we could identify genes that confer resistance against aminoglycosides, carbapenems, cephalosporins, glycylcyclines, lincosamides, macrolides, oxazolidinone, penams, phenicols, pleuromutilins, streptogramins, sulfonamides, tetracyclines, while other antibiotic groups including fluoroquinolones appeared in the genome of bacteria with a relatively lower clinical relevance, including *Escherichia coli*.

Such a great number and broad spectrum of ARGs and potentially affected antibiotic groups associated with the canine saliva samples may be related to the use of antibiotics at the small animal veterinary practice. Antibiotic consumption rates in companion animal sector are rather difficult to evaluate. However, some systems exist for the surveillance of magnitude of companion animal antibiotic consumption, such as European Surveillance of Veterinary Antimicrobial Consumption (ESVAC)^38^, VetCompass^39^ or the Small Animal Veterinary Surveillance Network (SAVSNET)^40^, these rates are still less well documented. Moreover, in many countries antimicrobial use is often just estimated of rough sales data^22^. Nevertheless, according to the two UK-based surveillance systems (VetCompass from Royal Veterinary College, and SAVSNET from Liverpool University) and one EU report (ESVAC) antibiotics are rather frequently prescribed at small animal clinics. A study states, 1 in 4 UK dogs (25.2%, 95% CI: 25.1–25.3%) was treated with antibiotics in a two-year period^41^. Even though, the vast majority of veterinarians are aware of the fact that improper AMU contributed to selection for antimicrobial resistance and that AMR is a significant problem according to nationwide surveys^42,43^, there are many factors that influence the antibiotics prescription preferences of veterinarians besides the perspectives of antimicrobial stewardship. According to a study conducted in Australia, veterinarians often report client pressure to prescribe antibiotics as the most significant factor limiting antibiotic stewardship goals^44^. In middle income countries, like South Africa cost may also influence the choice of antibiotics^45^. In contrast, in studies conducted by other research groups from countries including the U.S.^46^, the U.K.^47^, the Netherlands^48^ or Australia^49^, clinical settings were ranked of a much higher importance than client expectations. Broad-spectrum amoxicillin-clavulanate is the flagship of antimicrobial agents applied in dogs in many countries, while first-generation cephalosporins are also routinely used^22,50,51^. Lincosamides (clindamycin), macrolides, tetracyclines (doxycycline), nitroimidazoles and trimethoprim/sulphonamides have also been reported to be frequently used in small animal practice^22^. Third and fourth generation cephalosoprines, fluoroquinolones and polymixins that belong to cathegory B, ‘last resort’, or highest-priority Critically Important Antibiotics (HPCIAs) according to the European Medicines Agency^38^ should be avoided unless sensitivity testing is conducted, and no other antibiotics would be effective. Nevertheless, a HPCIAs have been estimated to be prescribed in around 5–6% of total antibiotic usage events. Of the HPCIA category, fluoroquinolones are the most common in dogs, constituting ~4 to 5% of total antibiotic prescriptions^52^.

In the current literature, human infections associated to dog bites are better- and more frequently documented than the transmission route of licking. Three to 30% of dog bites leads to infection^13^. Management of animal bites rests on two pillars: local wound care and adequately applied systemic treatment. Essentials of local therapy include inspection, debridement of the wound accompanied by the removal of possible foreign bodies, e.g. teeth and irrigation with saline solution. Additional radiologic diagnostics should be performed to rule out fracture and if clinically indicated. Recommendations on primary or delayed closure of the wound and analyses on risk of appearance of consequent infections are controversial since studies show that at least 6–8% of mammalian bites will become infected after primary closure. On the other hand, according to one randomized trial, facial wounds for example, have a low risk of infection due to their excellent blood supply even after primary closure. without the use of prophylactic antibiotics^53,54^. Wounds with delayed presentation and on the extremities should be left open^55^. Culturing of fresh bite wounds without signs of an abscess, severe cellulitis or sepsis can be avoided^56^. As for the systematic therapy, tetanus booster (if none given in past year) and rabies prophylaxis should always be considered. In our study, genome fragments of *Clostridium tetani*, the causative agent of tetanus was not detected in any of the examined saliva samples. No consensus has yet been found in the use of antibiotics for animal bite wound care. Prophylactic antibiotics should be considered unless the wound is very superficial and clean. Explicit indications for antibiotic prophylaxis or therapy include presentation at least 8 hours after the bite, clear signs of superinfection, moderate or severe wounds with crush injuries or devitalized tissues requiring surgery, deep puncture wounds (exceeding the layer of epidermis), wounds close to joints, diabetes mellitus, asplenic or immunocompromised state, alcohol abuse, or involvement of the genital area, face or hand^56–60^. In the absence of the above reasons, antibiotic therapy may not be necessary. Interestingly, injuries are normally located on the head, neck and face by children and on the hand or upper extremity by adults due to height ratios with the attacking dog^13,61^. The adequately chosen antibiotic agent is expected to be effective against anaerobe bacteria (*Bacteroides* spp., *Fusobacterium* spp., *Porphyromonas* spp., *Prevotella* spp. etc.), moreover *Staphylococcus, Streptococcus* and *Pasterurella* species. Prophylactic treatment is normally 3 to 5 days long, while medication for 10 days or longer is recommended if the wound is infected. The first-line choice for oral therapy is amoxicilin-clavulanate, accompanied with a first dose of intravenous antibiotic (e.g. ampicillinsulbactam, ticarcillin-clavulanate, piperacillin-tazobactam, or a carbapenem) in high risk patients. Amoxicillin-clavunalate is often combined with metronidazole or clindamycin, and is also sometimes replaced with cephalosporins, e.g. cefuroxime, cefotaxime, ceftriaxone, amoxicillin, fluoroquinolones, sulfamethoxazole and trimethoprim or, although less effective, azithromycin or doxycycline in this combination^56,60^. Due to high resistance rates, flucloxacillin, erythromycin and cephalosporins are often ineffective in *Pasteurella* infections, thus should rather be avoided^58^. In our case, no genes conferring resistance to these agent groups could be identified in *Pasturella* spp.

Data on the outcome of antibiotic prophylaxis in animal bite management by humans is limited and rather controversial and conflicting. While a meta-analysis of 8 randomized trial indicated a benefit of antibiotic prophylaxis^62^, some studies concluded that antibiotic prophylaxis does not result in a statistically significant difference in the frequency of wound infections among treated and untreated patient groups, except for the wounds to the hand^63^. Based on other publications, antibiotic prophylaxis should be recommended for the high-risk patient groups only^64,65^.

Amoxicillin-clavulanate, the most commonly used antibiotic in small animal medicine and the first choice for canine bite wounds is a member of broad-spectrum penicillins, that has been a frequently consumed key antibiotic group in the high-income super-region between 2000 and 2018 by a global study^66^. All in all, 7 ARG types were detected in the dog saliva samples that may confer resistance against amoxicillin-clavulanate, that were either the members of *blaTEM* or *OXA* family^67,68^. *TEM-116* was identified in *E. coli*, while various members of the *OXA* family appeared in in many genera, including *Acinetobacter baumannii, Bacteroides* spp., *Capnocytophaga* spp., *Fusobacterium ulcerans* and *Pseudomonas* spp. that can have a high clinical relevance in dog bite infections.

Many of the identified resistance genes harbored on iMGEs, phages or plasmids, with plasmids having the highest ARG association rates. Some genes could have been attached to two of the above mentioned mobility groups in the genome of one species, including the iMGE and phage co-appearance of aminoglycoside resistance encoding *aad(6)* and *aph(3’)-IIIa* in *Enterococcus faecium*, the iMGE and plasmid co-appearance of *aph(3’)-Ia* in *Corynebacterium* sp. 1959 and *Klebsiella quasipneumoniae, aph(3’)-Ia* and *aph(6)-Id* in *Variovorax* sp. SRS16, *aph(3”)-Ib* in *Variovorax* sp. PAMC28562, tetracycline resistance encoding *tetM* in *Enterococcus faecalis*, phage and plasmid co-appearance of macrolide, lincosamide and streptogramin resistance encoding *ermB* in *Enterococcus* sp. FDAARGOS_375. *OXA-2, OXA-22, OXA-347* and *TEM-116*, genes associated with amoxicillin-clavulanate resistance all appeared in plasmids in various species, moreover *OXA-2* was associated with both an iMGE and a plasmid in the genome of *Pseudomonas aeruginosa*. The accumulation of various mobility factors around the genes may increase the chance of the horizontal transfer of the given ARG. The canine saliva-borne transmission of bacteria harboring mobile ARGs may hamper antibiotic use in human clinical settings and can also contribute to the spread of AMR among the bacteria deriving from pets to the bacteriota appearing in humans.

In the contrary, canine saliva had been used to promote rapid healing and to reduce bacterial contamination in the past according to reports of etnoveterinary and ethnomedicinal practices^69,70^. Antimicrobial and anti-imflammatory activity of canine saliva induced by thiocyanate, lysozyme and indirectly, nitrate, among others^71,72^ can even appear at low concentrations^73^. However, according to our findings canine saliva can also be associated with public health risks, since salival bacteria may contaminate the surroundings of people and may also colonize human skin and mucous membranes. Thus ARG-rich bacteria present in and around humans do not even necessarily need to transfer their ARGs, to potentially cause severe harm to various groups of people with weaknesses of the immune system, e.g. extremities in age or diseased state.

As a common trend among many nations, veterinary use of antibiotics is gradually declining^38,44,52,74,75^. In human medicine, antibiotic sales elevated by 65% in low- and middle-income countries and decreased slightly, by 4% in high-income countries between 2000 and 2015, what adds up as a rise in global antibiotic consumption rates^66,76^. As a presumable conclusion, several genes conferring resistance against clinically important antibiotic groups are present in the salivary bacteriome of dogs that may drift to the genome of bacteria in humans. Encounters with dog saliva and dog bites may serve as an interspecies platform for the migration of bacteria and antimicrobial resistance genes. Transmitted bacteria may cause clinical symptoms and ARGs that they harbor may confer resistance against antibiotic agents of a clinical relevance.

## Declarations

### Ethics approval and consent to participate

Not applicable.

### Consent for publication

Not applicable.

### Availability of data and material

The datasets analysed in the current study are available in the National Center for Biotechnology Information (NCBI) Sequence Read Archive (SRA) repository and can be accessed through the PRJNA648123^77^ and PRJNA683923 BioProject identifiers.

### Author contributions statement

NS takes responsibility for the integrity of the data and the accuracy of the data analysis. AGT, LM and SN conceived the concept of the study. AGT, EGK and NS participated in the bioinformatic analysis. AGT, BR, EGK, IT and NS participated in the drafting of the manuscript. AD, AGT, ÁVP, BR, EGK, IT, LM, NS, SK and TN carried out the critical revision of the manuscript for important intellectual content. All authors read and approved the final manuscript.

## Acknowledgements

The authors would like to thank the providers of BioProject PRJNA648123^77^ and, furthermore, Kathleen Morrill; Elinor Karlsson, Ph.D.; Kerstin Lindblad-Toh, Ph.D. (Karlsson Laboratory, Broad Institute) and the Darwin’s Ark project (darwinsark.org) for providing the datasets and metadata for BioProject PRJNA683923.

## Authors’ information

No relevant information can be provided of the authors that may aid the readers’ interpretation of the article.

